# Cell wall inhibitors increase the accumulation of rifampicin in *Mycobacterium tuberculosis*

**DOI:** 10.1101/424309

**Authors:** Matthew B. McNeil, Somsundaram Chettiar, Divya Awasthi, Tanya Parish

## Abstract

There is a need for new combination regimens for tuberculosis. Identifying synergistic drug combinations can avoid toxic side effects and reduce treatment times. Using a fluorescent rifampicin conjugate, we demonstrated that synergy between cell wall inhibitors and rifampicin was associated with increased accumulation of rifampicin. Increased accumulation was also associated with increased cellular permeability.

---

*Mycobacterium tuberculosis* remains a major public health problem, causing approximately 1.7 million deaths in 2016 [1]. Current treatment regimens for *M. tuberculosis* involves a combination of 4 drugs, isoniazid (INH), rifampicin (RIF), pyrazinamide and ethambutol. Combination therapy is required to target *M. tuberculosis* in different physiological states. The emergence of resistance against existing drugs emphasizes the need for new therapeutic agents and combination regimens. Synergistic, additive or antagonistic drug interactions occur when the therapeutic activity of drug combinations are greater than, equal to, or less than the sum of the effects of the individual drugs [2]. Identifying synergy between existing TB drugs, or those late in development, would allow for new combination regimens that may reduce toxic side effects and the length of treatment.

MmpL3 transports trehalose monomycolates across the cytoplasmic membrane of *M. tuberculosis* and is the target of several families of inhibitors [3-6]. The MmpL3 inhibitor AU1235 has synergy with other TB agents including RIF, bedaquiline (BDQ) and the β-lactam ampicillin, with a reported fractional inhibitory index (FIC) of 0.5 when used in combination with all three agents [4]. Other MmpL3 inhibitors including the indole-2-carbozamides and SQ109 (FIC: 0.09) have synergy with RIF *in vitro* and in murine infection models [5, 7]. Synergy with RIF *in vitro* is also observed with other cell wall inhibitors including ethambutol and inhibitors of Pks13 (FIC: 0.55) [8, 9]. As the target of these inhibitors is the outer membrane of the cell wall, it is likely that the damaged mycolate layer would lead to increased cellular permeability resulting in increased accumulation and synergy with RIF. We wanted to test the hypothesis that synergy with RIF is due to increased permeability and intracellular accumulation.

We synthesized a fluorescent derivative or RIF by linking it to fluorescein isothiocyanate (RIF-FITC) (Figure 1) as described in the supplemental methods. RIF-FITC was active against *M. tuberculosis*, with a minimal inhibitory concentration (MIC) of 0.054 µM measured as described in [10, 11], although this is less active than the parent RIF molecule (MIC = 0.0060 µM). We confirmed that RIF-FITC was stable and did not undergo hydrolysis in PBS plus 0.05% w/v Tween (PBS-Tw), at 37°C for 3 h. We monitored the accumulation of RIF-FITC in wild-type *M. tuberculosis*. Bacteria were cultured in Middlebrook 7H9 medium plus 10% OADC supplement (oleic acid, albumin, destrose, catalase; Becton Dickinson) and 0.05 % w/v Tween 80 to an OD_590_ of 0.6-0.8, washed and resuspended in PBS-Tw to an OD_590_ of 0.6. RIF-FITC was added to cells (0.014-1.4 µM) and incubated at 37°C. Samples (3 mL) were harvested, washed in 10 mL PBS-Tw and resuspended in 2 mL PBS-Tw. Bacteria (100 µL) were dispensed into 96-well black-clear bottom plates and fluorescence measured at Ex490nm/Em525nm. We first confirmed that RIF-FITC accumulation could be detected. A starting concentration of 1.4 µM (25X MIC) RIF-FITC was required to produce a detectable signal (Figure 2a). Fluorescence increased in a linear fashion over 3 h (Figure 2a).

**Figure 1.**
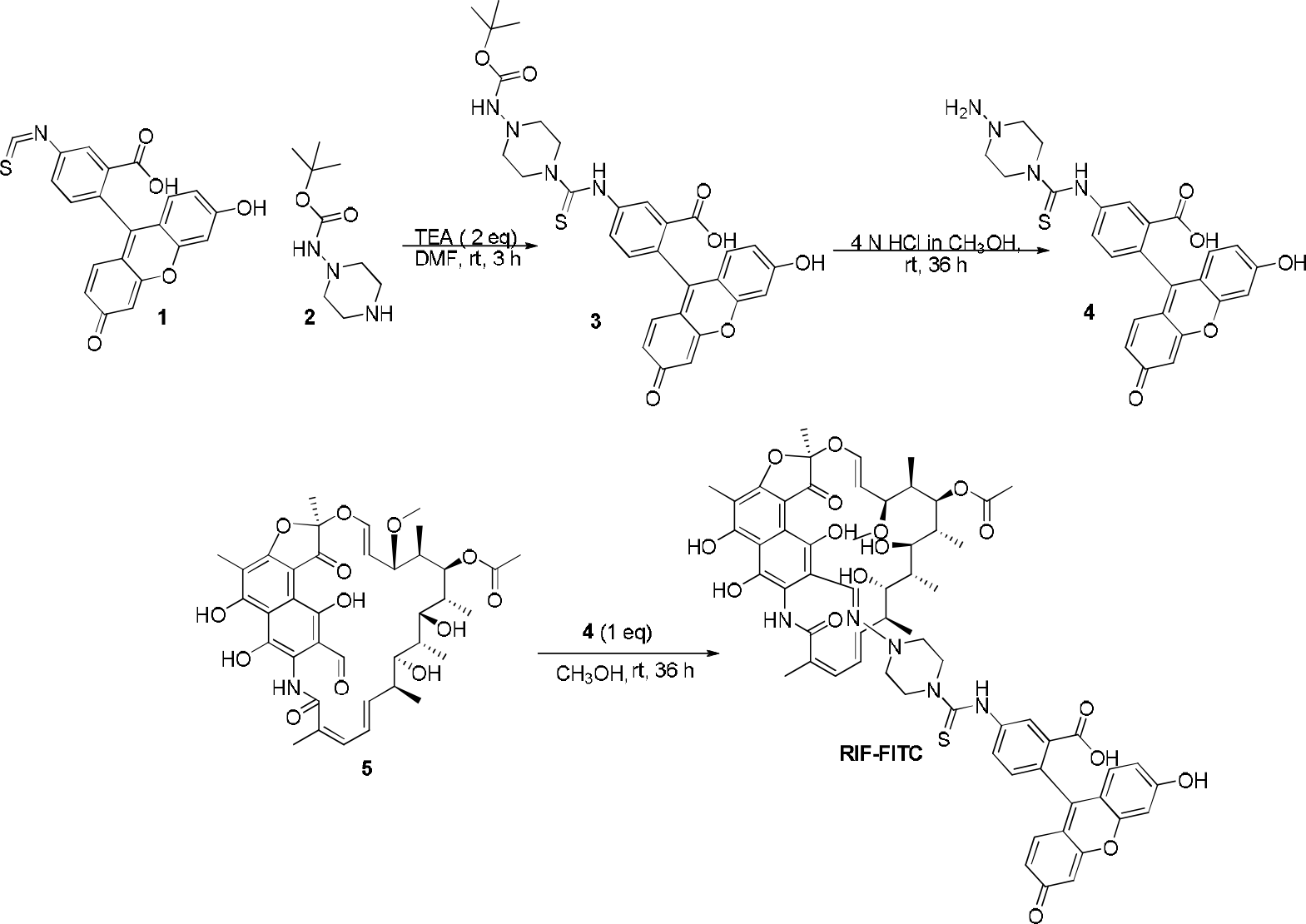
Synthesis of RIF-FITC: Schematic for the synthesis of RIF-FITC. TEA ̶ trimethylamine; DMF ̶ dimethylformamide.

**Figure 2.**
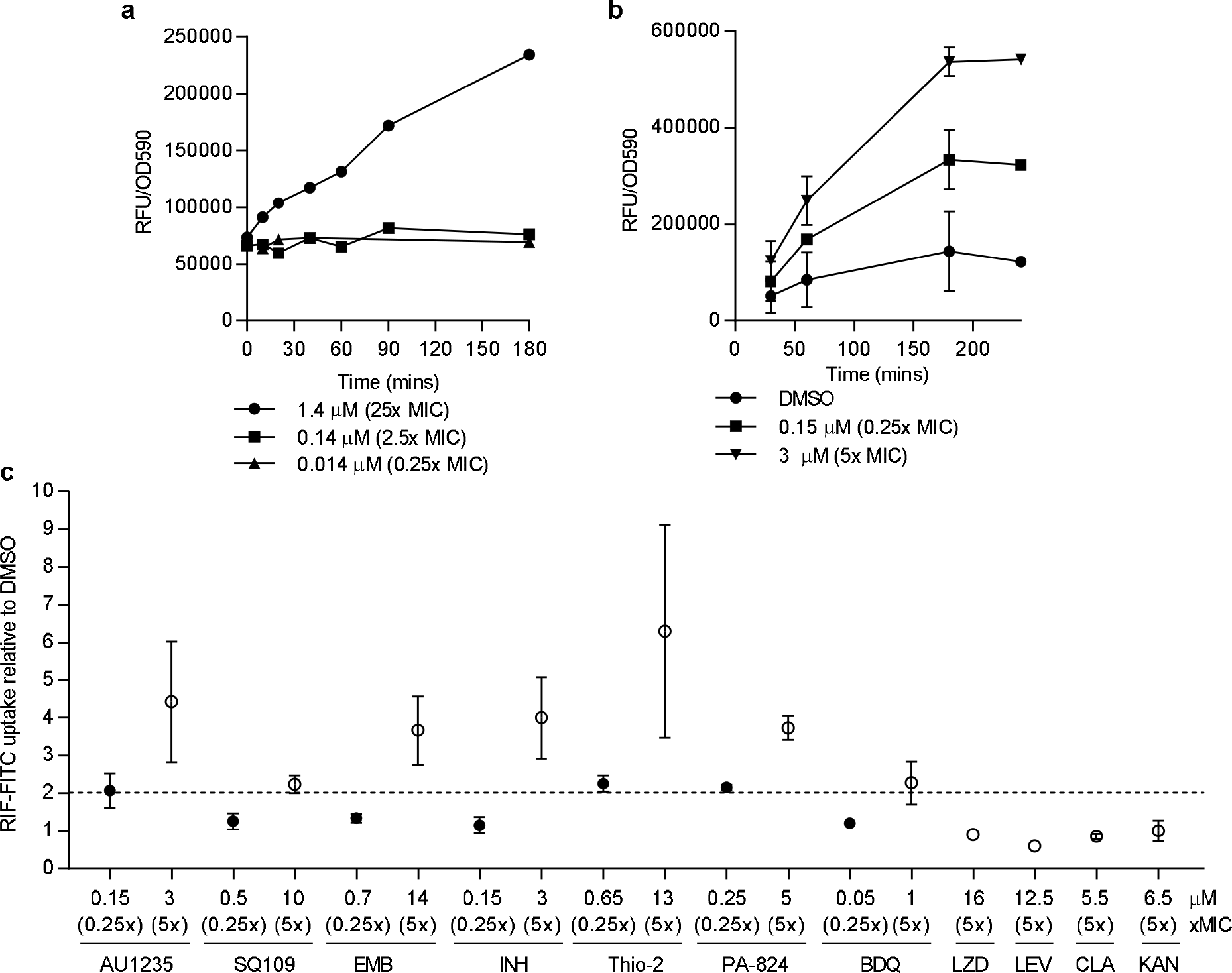
RIF-FITC accumulates in *M. tuberculosis*: (a) Accumulation of RIF-FITC into *M. tuberculosis*. (b) Accumulation of RIF-FITC in *M. tuberculosis* after pretreatment with 0.15 and 3 µM AU1235 for 24 h (n=2-3). (c) Accumulation of RIF-FITC in *M. tuberculosis* after treatment with compounds for 24 h (n= 2-5). Data are expressed as fold-change in RIF-FITC relative to the DMSO treated control. Results are the mean ± standard deviation. The dashed line represents a two-fold change relative to DMSO. MICs for compounds are AU1235 (0.6 µM), SQ109 (2 µM), EMB: Ethambutol (2.8 µM), INH: Isoniazid (0.6 µM), Thio-2: Thiophene-2 (2.6 µM), PA-824 (1 µM), BDQ: Bedaquiline (0.2 µM), LZD: Linezolid (3.2 µM), LEV: Levofloxacin (2.5 µM), CLA: Clarithromycin (1.1 µM), KAN: Kanamycin (1.3 µM).

The MmpL3 inhibitor AU1235 has synergy with RIF *in vitro*, with a reported FIC of 0.5 [4]. We determined whether treatment with AU1235 could affect RIF-FITC accumulation. *M. tuberculosis* was grown to an OD_590_ of 0.3, compound added and cells cultured for a further 24 h. RIF-FITC was added to cells and cultures incubated at 37°C. Samples (3 mL) were harvested, washed in 10 mL PBS-Tw and resuspended in 2 mL PBS-Tw. Bacteria (100 µL) were dispensed into 96-well black-clear bottom plates and fluorescence measured at Ex 490 nm, Em 525nm. The accumulation of RIF-FITC increased two-fold after 3 h in the presence of a sub-inhibitory concentration of AU1235 (0.15 µM or 0.25X MIC) (Figure 2b). At concentrations above the MIC (3 µM or 5X MIC), there was a four-fold increase in intracellular RIF-FITC accumulation (Figure 2b). Thus, we saw increased accumulation of RIF-FITC in *M. tuberculosis* following exposure to the MmpL3 inhibitor, AU1235. No aggregation was observed in the cutlures after addition of compounds.

We monitored the accumulation of RIF-FITC in *M. tuberculosis* following exposure to other anti-tubercular agents. We tested RIF-FITC accumulation after 3 h incubation, as we expected this to be at steady state as demonstrated in Figure 2b. At above inhibitory concentrations (i.e. 5x MIC) the cell wall inhibitors, AU1235 (MmpL3 inhibitor) [6], ethambutol (arabinosyl transferase inhibitor) [12], INH (InhA inhibitor) [13] and thiophene-2 (Pks13 inhibitor) [14] resulted in at least a four-fold increase in RIF-FITC accumulation (Figure 2c). An alternative MmpL3 inhibitor, SQ109, increased RIF-FITC accumulation two-fold (Figure 2c). The respiratory inhibitors PA-824 [15] and BDQ [16], increased RIF-FITC accumulation by four-fold and two-fold respectively (Figure 2c). At sub-inhibitory concentrations (0.25X MIC) AU1235, thiophene-2 and PA-824 all increased RIF-FITC accumulation by two-fold (Figure 2c). INH, ethambutol, SQ109 and BDQ failed to increase RIF-FITC accumulation at 0.25X MIC (Figure 2c). Other anti-tubercular agents that do not target the cell wall (linezolid, levofloxacin, clarithromycin and kanamycin) had no effect on RIF-FITC accumulation at 5X MIC (Figure 2c). In conclusion, a variety of cell wall and respiratory inhibitors increase the accumulation of RIF-FITC into *M. tuberculosis* and a subset of those inhibitors were able to increase RIF-FITC accumulation at sub-inhibitory concentrations.

To determine if increased accumulation of RIF-FITC was due to increased permeability, we used the ethidium bromide (EtBr) assay [17]. *M. tuberculosis* was grown to an OD_590_ of 0.3. Compounds were added and cultures grown for approximately 24 h; cells were harvested, washed and resuspended in PBS-Tw buffer to an OD_590_ of 0.8. An equal volume of culture was added to 50 µL of PBS-Tw containing 8 µg/mL EtBr in 96-well plates. Intracellular accumulation of EtBr was monitored at 37°C using Ex 530 nm, Em 590 nm. AU1235 and PA824, both of which increased RIF-FITC accumulation at 5X and 0.25X MIC increased EtBr accumulation at 5X and 0.25X MIC (Figure 3a and b). At 0.25X MIC the rate of accumulation was lower than 5X, although a similar steady state level was reached (Figure 3a and b). SQ109 and INH, which increased RIF-FITC accumulation at 5X MIC but not at 0.25X MIC increased the rate of accumulation and steady state levels of EtBr at 5X MIC but not at 0.25X MIC (Figure 3d and e). Ethambutol, which increased RIF-FITC accumulation at 5X but not at 0.25X MIC increased EtBr accumulation at both 5X and 0.25X (Figure 3f). This correlation suggests that the compounds are increasing cell wall permeability and/or disrupting efflux, allowing for greater intracellular accumulation of both RIF and EtBr. Interestingly, thiophene-2 and BDQ which increased RIF-FITC accumulation at 5X MIC resulted in reduced EtBr accumulation as compared to DMSO treated cells (Figure 3g and h). This suggests that the mechanisms for increased RIF accumulation for these two compounds are different and are not related simply to changes in cell wall permeability and/or efflux. Since BDQ disrupts ATP generation, it could have negative effects on ATP-dependent transport mechanisms which might be independent of changes in cell wall structure.

**Figure 3.**
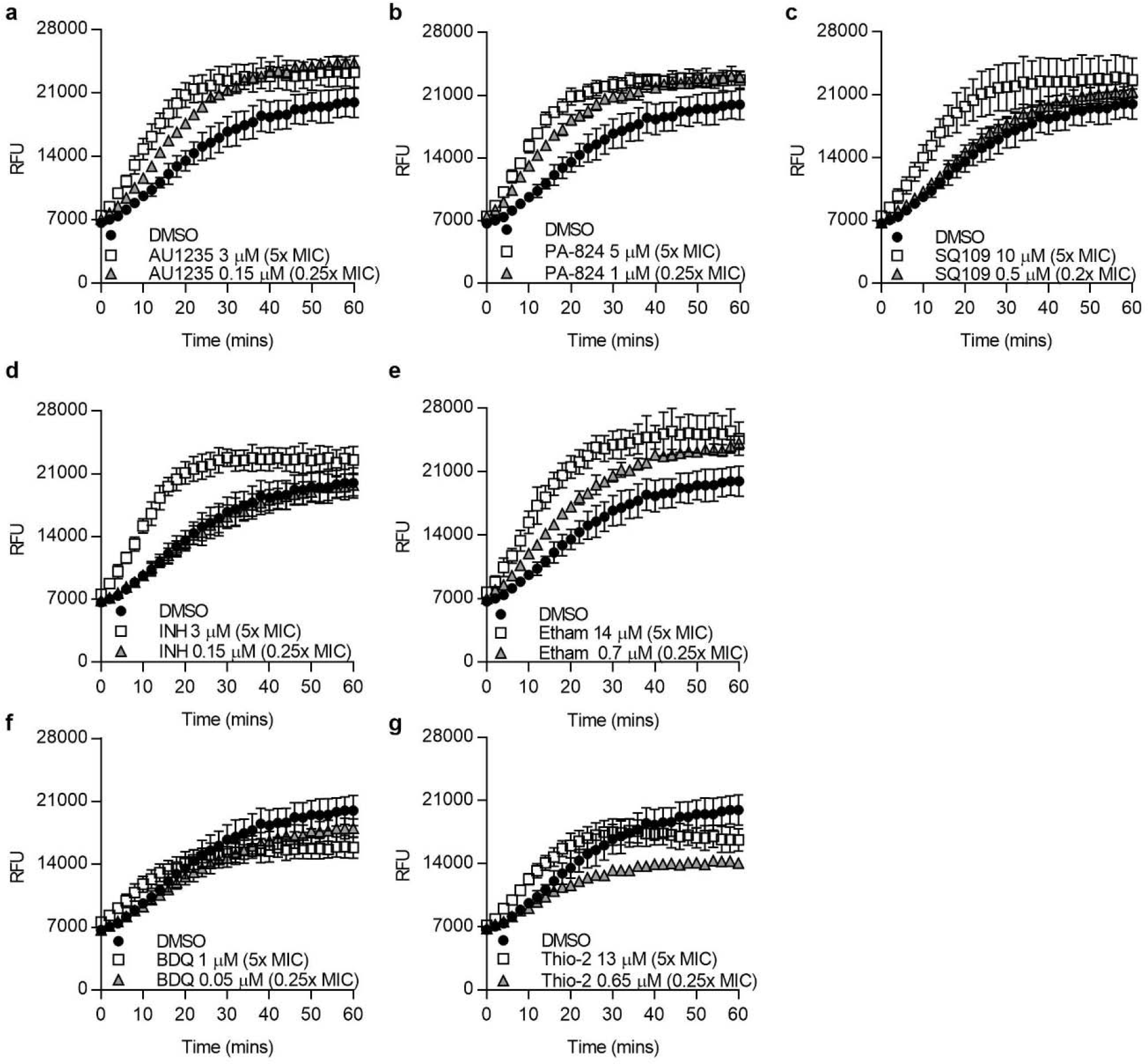
Cell wall inhibitors increase permeability of *M. tuberculosis*: *M. tuberculosis* was pretreated with compounds for 24 h. Ethidium bromide uptake was monitored by fluorescence (Ex530/Em590). (a) AU1235, (b) PA-824, (c) SQ109, (d) INH, (e) ethambutol, (f) BDQ and (g) Thiophene-2. Relative fluorescent units (RFU) from every second minute is presented. Results are mean ± standard deviation from technical and biological duplicates (n=4).

This current study demonstrates that synergistic interactions between RIF and compounds that target proteins involved in cell wall synthesis, including MmpL3 and Pks13, are consistent with disruptions in the cell wall leading to increased cellular permeability and increased accumulation of RIF. Increased accumulation of RIF in ethambutol-treated bacteria is consistent with previous studies that detected intracellular RIF using radio-labelling or liquid chromatography [8, 18]. The observation that INH did not increase RIF accumulation at sub-inhibitory concentrations is consistent with previous work showing a lack of synergy between the two agents in *Mycobacterium bovis* BCG [18]. However, we demonstrate that at concentrations above the MIC, INH is able to increase the accumulation of RIF. The respiratory inhibitor PA-824 has multiple modes of action, including disruptions in cell wall [15, 19]. We hypothesize, that PA-824 mediated disruptions of the cell wall are responsible for increased permeability and accumulation of RIF. Further studies are required to determine whether synergy between cell wall inhibitors and other compounds is due to increased accumulation and whether increased accumulation translates to improved activity when used in combination.

## Acknowledgments

The authors acknowledge Catherine Shelton for technical assistance.

